# Characterization of Copper Complex Nanoparticles synthesized by Plant Polyphenols

**DOI:** 10.1101/134940

**Authors:** Zhiqiang Wang, Pengfei An

## Abstract

In this paper a kind of copper oxide material of copper–polyphenols complex nanoparticles (Cu–P NPs) were synthesized by *Cinnamomum pedunculatum* leaves extract, which have different morphology and appearance compared with usual copper oxides Cu_2_O and CuO. For better understanding about this material, the Cu–P NPs were characterized using scanning electron microscopy (SEM), X-ray absorption spectroscopy (XAS), X-ray diffraction (XRD), Fourier transform infrared spectroscopy (FTIR), and thermogravimetric analysis (TGA). It was found that synthesized Cu–P NPs were amorphous with spherical particles ranged from 80 to 500 nm. XAS data analysis indicated that the synthesized Cu–P NPs has different molecular structure with Cu_2_O and CuO. It is assumed that the copper ions chelated with polyphenol molecule. The nanoparticles showed a clear anti *Escherichia coli* activity in this study, and may also be used in fields of semiconductor, ceramic, catalyst, and sensor. This synthesis approach provided a novel route to manufacture copper oxide nanomaterial.

## 1. Introduction

Copper oxide nanoparticles can be used as catalysts for reduction, oxidation, electrocatalysis, photocatalysis, and gas-phase reaction [1]. Various physical and chemical methods have been extensively used to produce copper oxide nanoparticles such as chemical treatment, microemulsion method, arc-submerged nanoparticle synthesis system, flame-based aerosol methods, sonochemcial, hydrothermal and solid-state techniques [2-5]. Chemical treatment is regarded more usual methods providing advantages in terms of shape and/or size selectivity. These methods apply toxic chemicals or require high energy usually. Therefore, development of clean, biocompatible, eco-friendly methods for nanoparticles synthesis deserve merit. Review of literature revealed that there are only a very few literatures reported on the use of yeast, fungi, bacteria or plant extract for synthesizing copper oxide NPs [6-10]. However, there is still some ambiguity about the binding structures between organic matter and chelated Cu^2+^ ions. In 2013, our study [11] reported iron–polyphenols NPs’ molecular structure by using XAS analysis, revealed that trivalent iron ions can chelated with oxygen atoms of plant polyphenols to form nanoparticles. Therefore, we would like to investigate copper– polyphenols NPs using XAS again to determine their molecular structure at atom level. And also some studies of the coordination environment of Cu^2+^ ions in a mixture with polyphenols, probed using SEM, XRD, FTIR, and TGA have been done. It was found the mixture of copper salt solution with plant leaves extract leads to formation of copper– polyphenol complex nanoparticles, whose appearance and characters are different from either the reagent copper sulfate or conventional copper oxides.

## 2. Material and methods

### Synthesis of Copper – Polyphenol Complex NPs

The plant leaves extract was prepared with the method we used before [12], by heating 200 g *Cinnamomum pedunculatum* leaves (collected from Nanshan botanical garden Chongqing, China) added to 1000 mL Milli–Q water at 60 °C for 2 h. After settling for 0.5 h, the extract was vacuum-filtered. A solution of 0.10 M CuSO_4_ was prepared by adding 250 g of CuSO_4_•5H_2_O (Sigma– Aldrich) in 1.0 L of Milli–Q water, and subsequently the leaf extract was added to 0.1 M CuSO_4_ solution in 1:1 ratio in volume. The formation of copper containing NPs was marked by the appearance of a yellowish brown colour. The NPs were separated by centrifuging at 4000 rpm and dried in open air for 24 hours.

### Characterizations and Measurements

#### Scanning Electron Microscopy

Morphological characteristics were analysed using a Field Emission-Scanning electron microscope (FE-SEM), specifically an FEI Instruments-Quanta-450-FEG.

#### XAS Data Collection

X-ray absorption spectroscopy (XAS) experiments were performed at the Beijing Synchrotron Radiation facility (BSRF). The storage ring operates at 1.5 GeV and a typical current of 200 mA. A Si(111) double crystal was employed as a monochromator. Data were acquired in transmission mode. The ionization chamber was filled with an N_2_ gas.

#### XAS Data analysis

XANES spectra were normalized to an edge jump of unity taking into account the atomic background after the edge as it comes out from the EXAFS analysis. Prior removal of the background absorption was done by subtraction of a linear function extrapolated from the pre-edge region. XAS analysis has been performed by using the Athena and Artemis package [13] that takes into account the multiple scattering theory. The method allows the direct comparison of the raw experimental data with a model theoretical signal.

#### X–ray Diffraction

XRD patterns of Cu*–*P NPs samples were obtained using a Philips PANalytical-Empyrean instrument. The source consisted of Cu Kα radiation (λ = 1.54 Å). Each sample was dried in an open area for 24 h and then scanned in the 2θ range of 10*–* 80 °.

#### Fourier Transform Infra-red Spectroscopy

Agilent 660-IR was used to examine the functioning groups on the Cu–P NPs’ surface. Samples for FTIR measurement were prepared by mixing 1% (w/w) specimen with 100 mg of KBr powder and pressed into a sheer slice. An average of 32 scans was collected for each measurement employing a resolution of 2 cm.

#### Thermogravimetric Analysis (TGA)

The thermal stability and resistance to decomposition of the polymer were recorded on a thermogravimetric analyzer (SDT Q 600, T.A. Instruments-water LLC, Newcastle) in a temperature range of 50-900 °C under constant nitrogen flow (100 mL/min).

## 3. Results and Discussion

The Picture 1 shows the synthesized Cu–P NPs in paste state by centrifuging after synthesis and dry powder, which was exposed in air for 24 h. They have brown colour, which are different from the black colour of CuO, and red colour of Cu_2_O. It indicated the synthesized material may be not the CuO and Cu_2_O.

**Picture 1:**
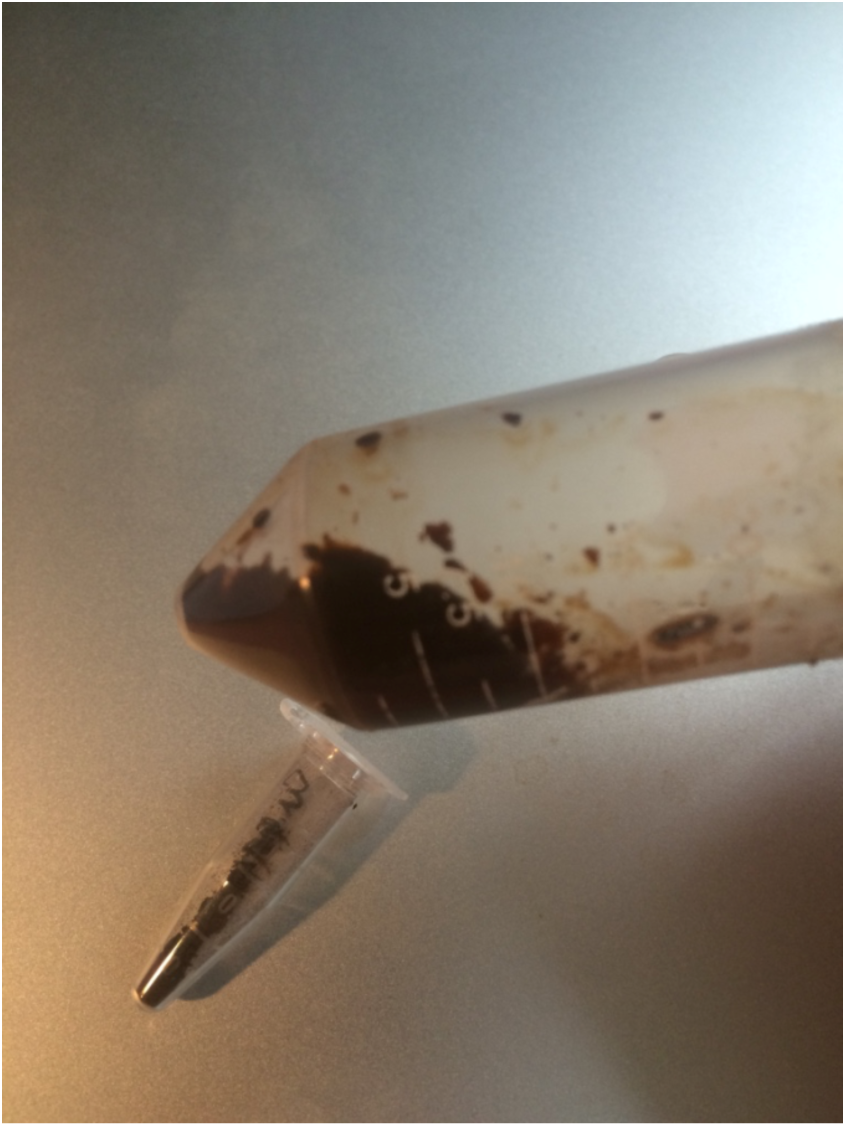
Synthesized Cu–P NPs in paste state (in the big centrifuge tube) and their dry powder (in the small centrifuge tube).

### SEM

In Figure 1 the SEM images revealed the status of Cu–P NPs in states of colloid. The NPs in colloid were in range of 80–900 nm spherical particles capped by plant organic matters. The Figure 2 image showed that the powder become crystals after exposure in the air for 24 h. According to our previous study [14] about iron–polyphenols NPs, their sizes and appearance can be determined by reagent concentration, type of polyphenols and temperature. This result could be also applied for copper–polyphenols NPs.

**Figure 1:**
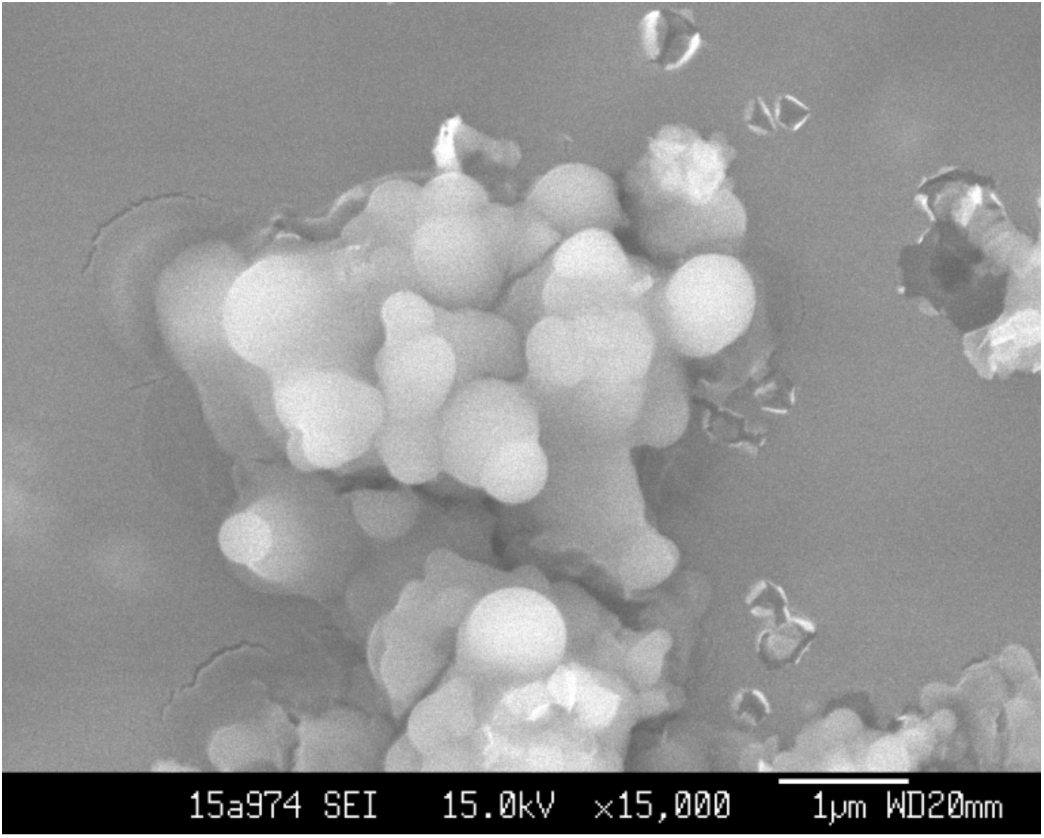
SEM image of Cu–P NPs in colloid.

**Figure 2:**
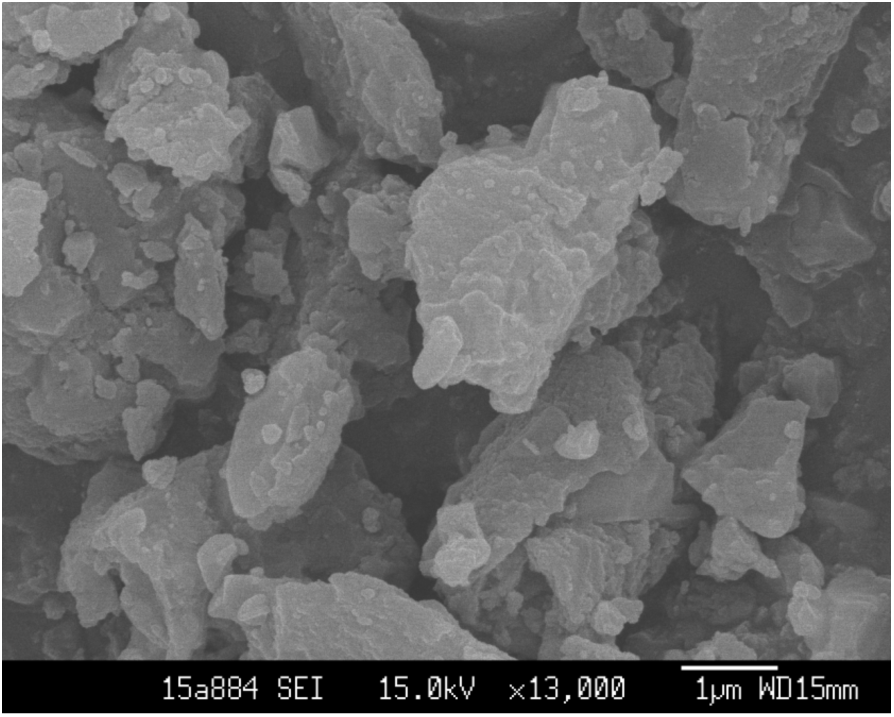
SEM image of dry Cu–P NPs powder.

### XANES and EXAFS Analysis

The Cu K-edge XANES analysis is used in this work to determine the average Cu valence state in the Cu-P NPs sample. The energy position of the Cu K-edge in Cu-P NPs sample is close to the edge position of the Cu^2+^ compounds (Figure 3), confirming that the copper in the sample is predominantly divalent ion. At the same time, we can clearly see that the curve of Cu–P NPs is none of the model samples, indicating that Cu–P NPs have different molecular structure from these model copper– containing materials.

**Figure 3:**
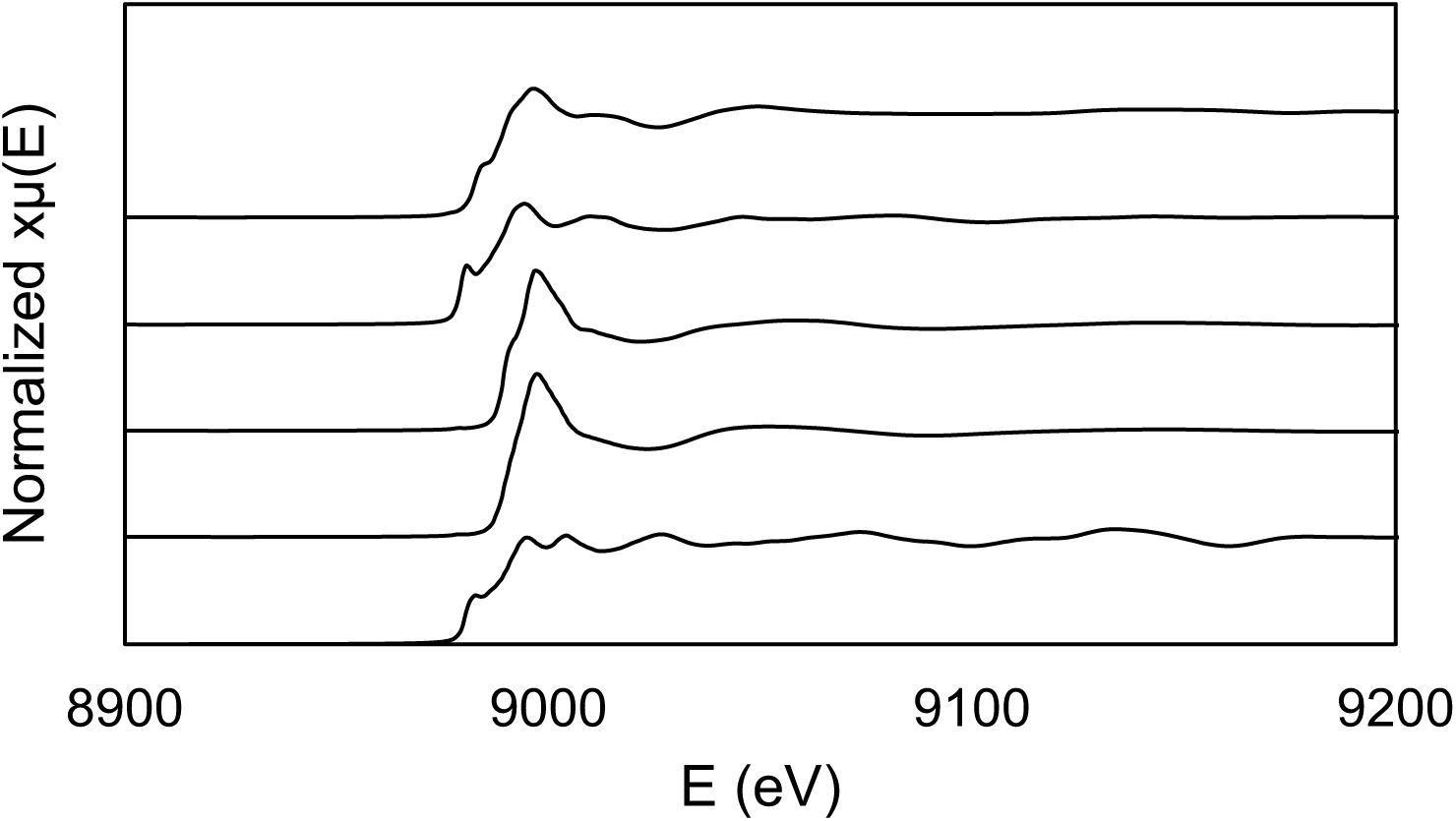
XANES spectrum of Cu–P NPs.

The EXAFS simulations suggested that Cu K-edge EXAFS analysis is used to directly probe the local structure around Cu cation in the Cu–P NPs. In the Fourier transform magnitude of the EXAFS spectrum (Figure 4), three distinct peaks are observed representing the contributions of photoelectron scattering on the nearest shells of neighbors around the Cu atom. Three variable parameters for each shell of neighbors are introduced in the model: the shell coordination number (*N*), the distance (*R*) and the Debye–Waller factor (*σ*^2^). A good agreement between the model and experimental spectrum is found using the *R* range from 0 to 5 Å. The list of best fit parameters is given in Table 1. One oxygen atom is identified in the fit of the Cu-O distance of 1.46 Å, which is different from the Cu-O distances of CuO and Cu_2_O as reported [15]. This result indicated that the synthesized material has different molecular structure with CuO and Cu_2_O. Relatively large Debye–Waller factors for the Cu coordination shell indicate large disorder in the structure. The XANES and EXAFS results indicated that the synthesized material is a kind of copper oxide, but has different molecular structure with Cu_2_O and CuO due to chelating bond formed between copper ion and oxygen atom from polyphenol molecule. Guo *et al.* [16] assumed that Cu^2+^ ions can react with tannic acid to form a phenolic network capsule with Energy-dispersive X-ray elemental mapping. Our research [17] revealed that the phenolic network containing metal ions, oxygen and carbon atoms takes place due to condensation of polyphenols when adding metal ions in them, which results in losing water molecule between polyphenol molecules. It is well known that chelation takes place between metal ions and organic matter. Our works [18] have proven that the chelation between metal ions and organic matter can form nanoparticles. We found that plant polyphenols can reduce gold and silver salts in solution to pure gold and silver nanoparticles, but can only chelate copper and other metal elements, which have higher reducing potentials [17]. Therefore, the synthesized material may be called copper-polyphenol chelating complex nanoparticles.

**Table 1:** Structural parameters *R* (Å), *CN* and *σ* (Å) derived from the EXAFS data analysis of the Fe-P NPs. (*R* – interatomic distance; *CN* – coordination number; *σ* – Debye-Waller factor)

| Bond | R(Å) | N | $\sigma^2(\text{\AA}^2)$ |
| --- | --- | --- | --- |
| Cu-O | 1.46 | 1 | 0.004 |

**Figure 4:**
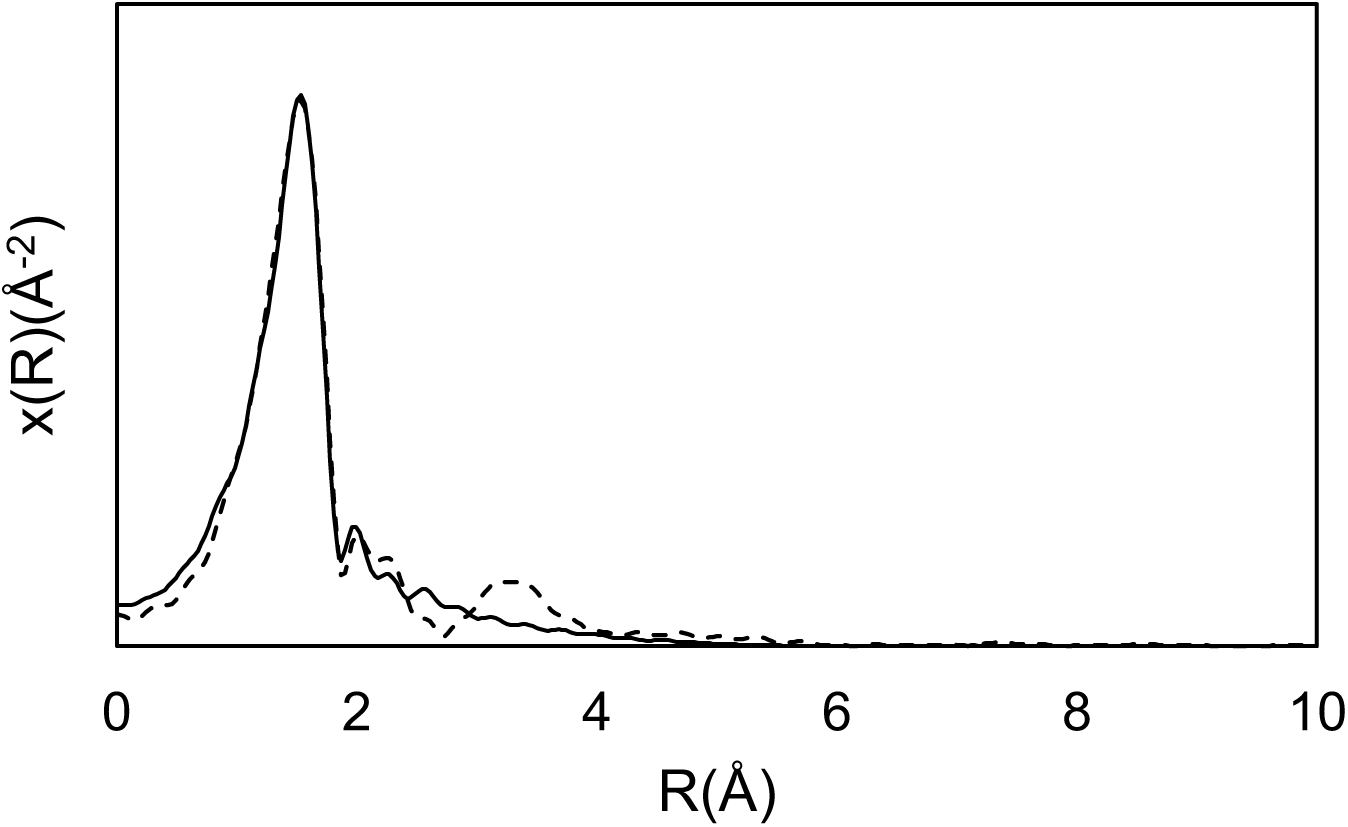
EXAFS spectrum of Cu–P NPs. Experimental (solid line); theoretical (dots).

### X–ray Diffraction

The XRD patterns of the Cu–P NPs paste did not show distinct diffraction peaks, suggesting that the material is amorphous (Figure 5). The dry powder showed typical crystal XRD patterns which don’t consist with any pure crystal material (Figure 6). This means the powder is heterogeneous material. It is assumed that plant polyphenols capped the regent CuSO_4_ solution by hydrogen bond, formed amorphous molecular structure, and changed the bond connections among Cu^2+^ and SO_4_^2-^ in some degree. After 24 h drying in air, the Cu–P NPs were oxidized and became crystals.

**Figure 5:**
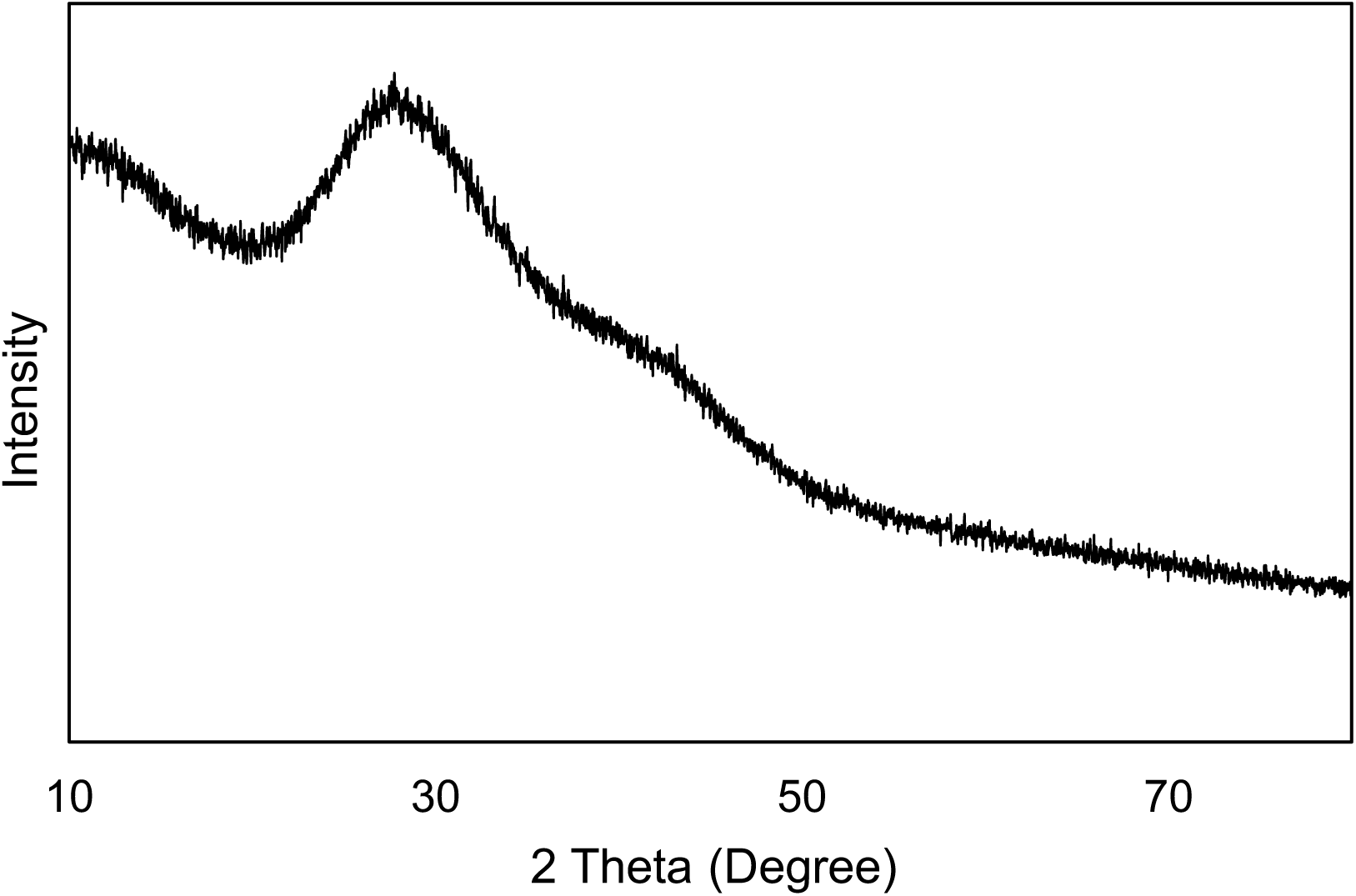
XRD spectrum of wet Cu–P NPs paste.

**Figure 6:**
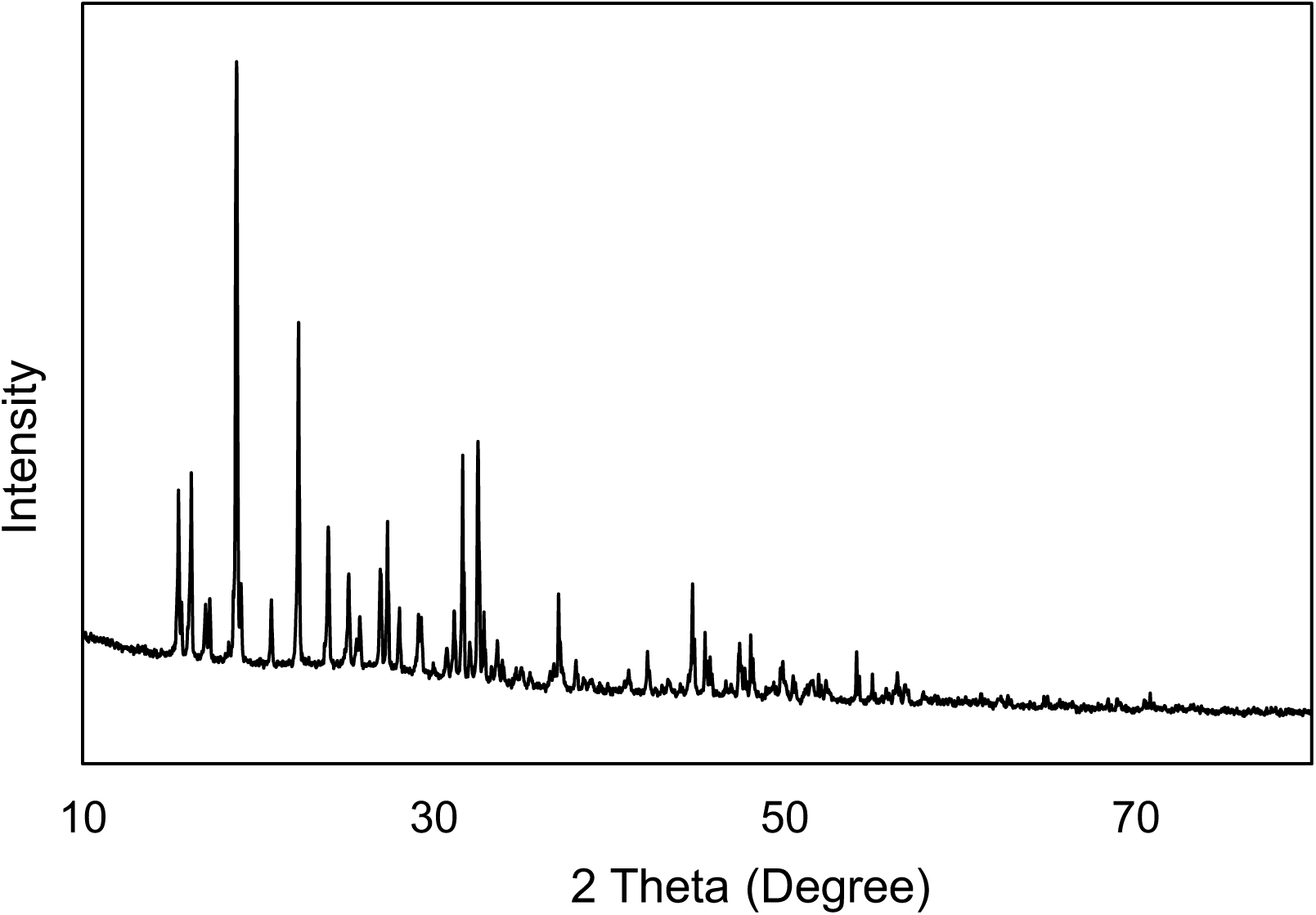
XRD spectrum of dry Cu–P NPs powder.

### Fourier Transform Infra-red Spectroscopy

Figure 7 illustrates the FTIR spectrum of the Cu–P NPs powder sample, revealing several peaks in the spectral range 1000–4000 cm^−1^. The peak at 3430 cm^−1^ is attributed to O–H of the H bond or carboxylic acid. The feature around 1640 cm^−1^ presumably corresponds to the O–H of carboxylic acids. The peak around 1200 cm^−1^ and 1160 cm^−1^ can be attributed to the C–O of carboxylic acid. All these features indicated this dry powder nanomaterial is capped by organic matter. This outcome agrees with the results of XAS and XRD investigations.

**Figure 7:**
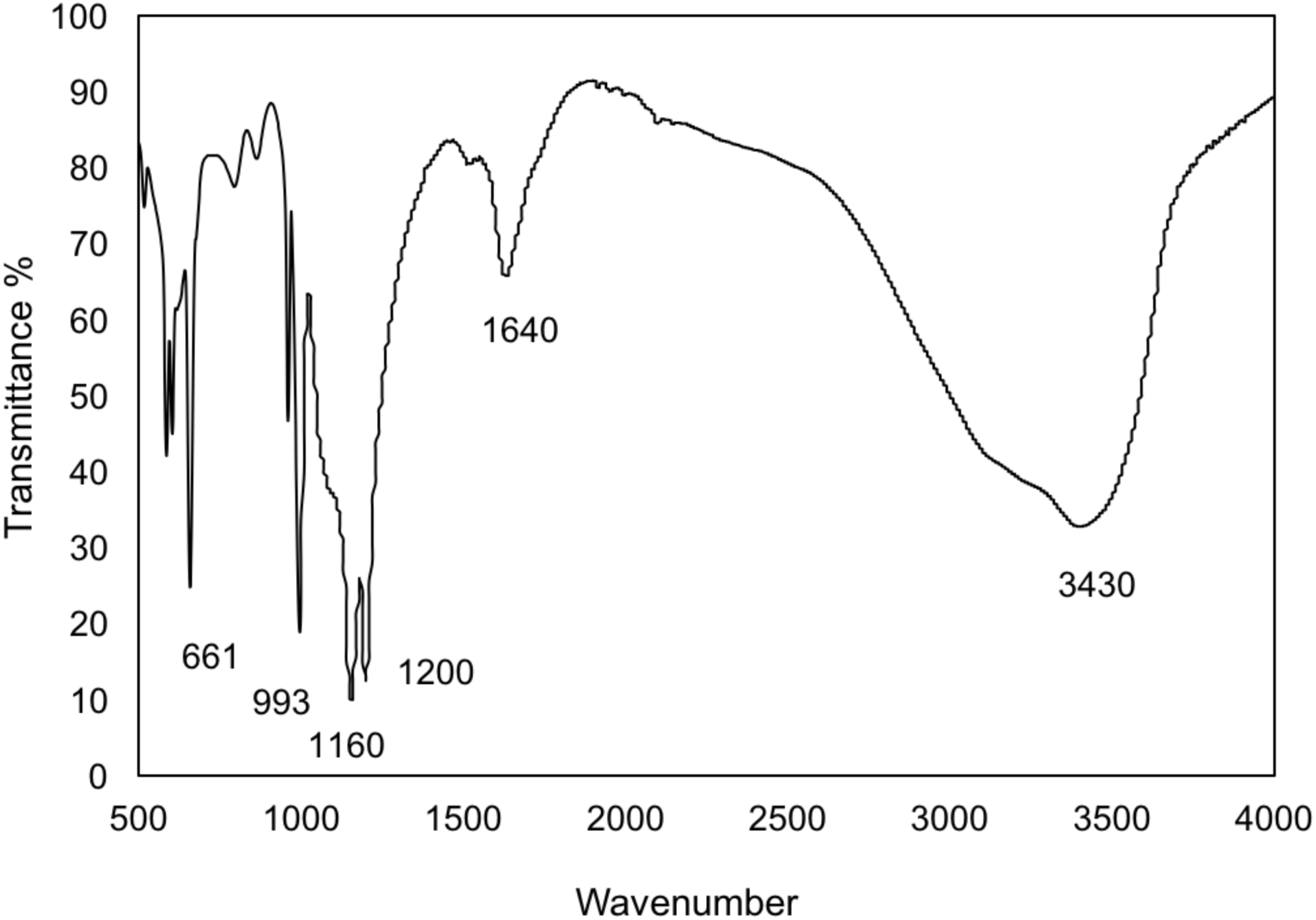
FTIR spectrum of Cu–P NPs.

**Figure 8:**
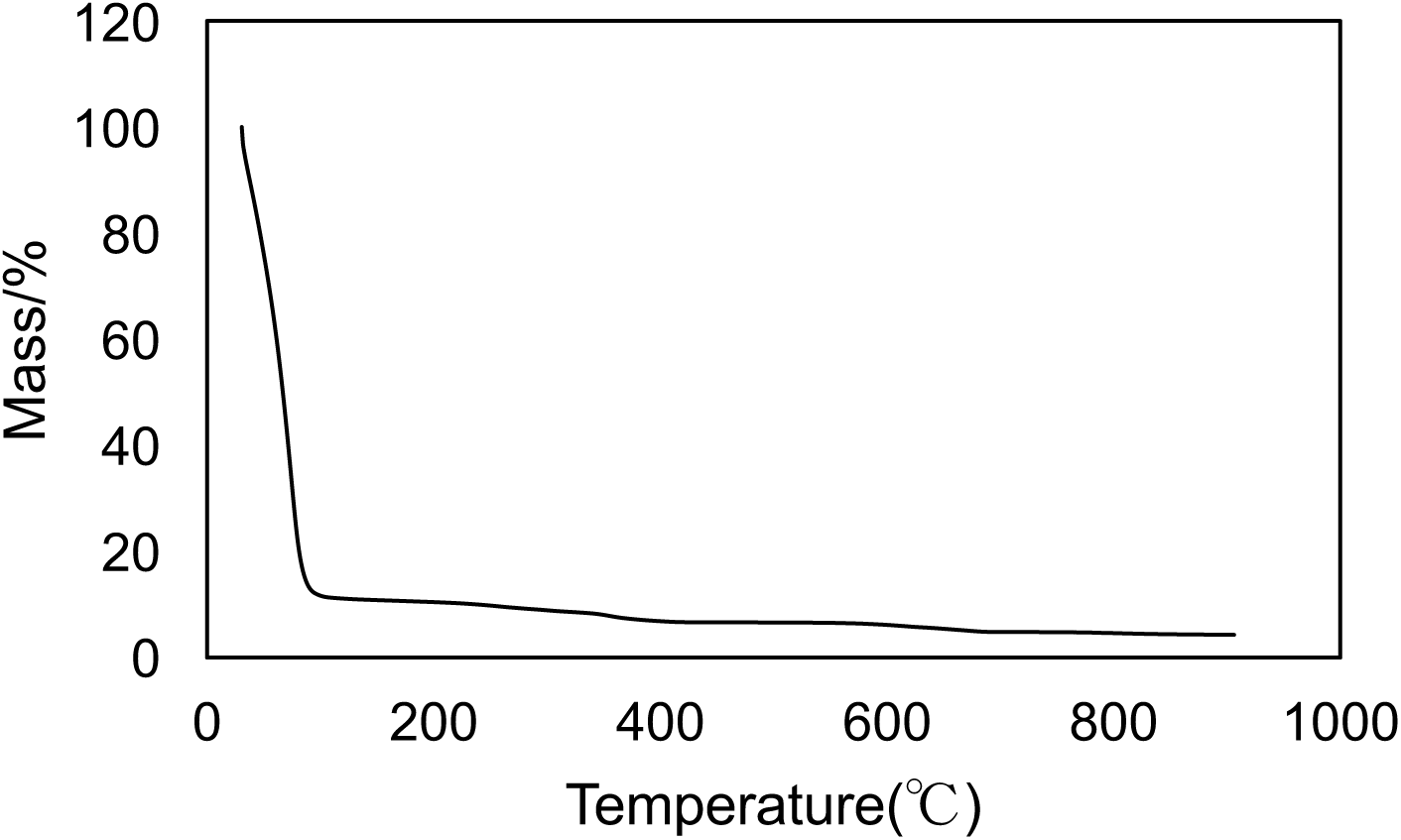
TG spectrum of Cu–P NPs.

### Thermogravimetric Analysis (TGA)

The thermal stability of the polycopper sulphate NPs was determined by thermogravimetric analysis. The TGA profile of the NPs indicated a two-step mass loss: almost 88% of total mass was lost in the temperature range up to 95 °C. This may be attributed to water loss due to the porous structure of the material, which can contain large amount of water. The mass percentage was reduced from 12% to 4.2% gradually with temperature increase from 96 to 900 °C. It is assumed that organic matter chelated with copper ions was removed with increasing temperature. The residue may be copper oxide. This phenomenon suggested that the synthesized nanoparticles have big surface area, which can adsorb large amount of water, organic matter or other substances.

### Antibacterial activity studies

The antibacterial activity of Cu–P NPs was carried out on human pathogenic *Escherichia coli*, by standard disc diffusion method. Mueller Hinton agar medium was used to cultivate bacteria. Standard antibiotic dis (100μg/ml) Ampicilin was used as reference drug and plant extract used as a control. A clear 100% inhibition zone has been found after adding of Cu–P NPs into the cultivated *Escherichia coli*. This means that the synthesized Cu–P NPs were highly effective in their activity against these bacteria, because the released copper ions from the Cu–P NPs the cause the membrane to rupture and kill bacteria.

## 4. Conclusions

The characterization of Cu–P NPs using *Cinnamomum pedunculatum* leaves has been demonstrated. The XANES and EXAFS results indicated that the synthesized material is a kind of copper oxide, but has different molecular structure with its reagent CuSO_4_ and the conventional copper oxides such as Cu_2_O and CuO. The XRD and FTIR results suggested that the synthesized materials are chelated and capped with plant polyphenols. Therefore, it can be supposed that heterogeneous copper–polyphenols complex nanoparticles were formed. TGA pattern showed that the Cu–P NPs contain 88% water and up to 7.8 % organic matter in weight. The nanoparticles showed a clear anti *Escherichia coli* activity. This green synthesis approach provided a route to synthesis copper–polyphenol complex nanoparticles, which may serve as nanoagent in other applications such as semiconductor, medicine, catalyst, ceramic, and sensors.

